# Prolonged mitosis causes separase deregulation and chromosome nondisjunction

**DOI:** 10.1101/2020.06.30.179267

**Authors:** Norihisa Shindo, Makoto Otsuki, Kazuhiko S.K. Uchida, Toru Hirota

**Affiliations:** Division of Experimental Pathology, Cancer Institute of the Japanese Foundation for Cancer Research (JFCR), Ariake 3-8-31 Koto-ku, 135-8550 Tokyo, Japan

## Abstract

Intercellular karyotype diversity, or aneuploidy, is a widespread feature of cancers propagated through continuous mitotic chromosome segregation errors, the condition called chromosomal instability. The protease separase is an essential enzyme that sever cohesin between sister chromatids, and a probe for separase activity has articulated that separase undergoes abrupt activation shortly before anaphase onset, after being suppressed through metaphase; however the relevance of this robust control remains unclear. In this study, we found that separase activates precociously during prolonged metaphase, consistently in multiple types of cancer cell lines. An artificial extension of metaphase alone in chromosomally stable diploid cells was found to induce precocious activation. These kinetic changes resulted in an incomplete removal of cohesin and emergence of chromosomal bridges in anaphase. Conversely, in transformed cells, shortening back of their prolonged metaphase restored the robust activation of separase and ameliorated anaphase bridge formation. These observations suggest that retarded metaphase progression directly affects the separase activation profile, which provides a previously unanticipated etiology for chromosomal instability in cancers and underscores the significance of the swift mitotic transitions for fail-safe chromosome segregation.

## Introduction

Aneuploidy is a widespread feature of malignancies with advanced disease that is generated by persistent chromosome segregation failure as cells divide. This pathological condition called chromosome instability can be manifested as lagging and bridging chromosomes in anaphase. Lagging chromosomes largely accounts for numerical aberrations and chromosomal bridges structural aberrations, both of which comprises intercellular karyotype diversity in cancers (Reviewed in Thompson et al., 2010).

Lagging chromosomes is known to arise primarily from error-prone kinetochore-microtubule interactions, namely merotelic attachments. The merotelic attachments are normally dissolved during early mitosis through the mechanisms requiring dynamics of kinetochore microtubules (Bakhoum et al., 2009a) and proficient activity of Aurora B (Cimini et al., 2006). Accordingly, loss of microtubule dynamics (Bakhoum et al., 2009b) and insufficient centromeric activity of Aurora B (Abe et al., 2016) are the features shared among various types of cancer cells, explaining their elevated incidence of lagging chromosomes. By contrast, causes of chromosomal bridges are thought to relate more to chromosomal problem(s) outside mitosis, including defective chromatid assembly, incomplete replication, and dicentric chromosomes (Mankouri et al., 2013; Ganem and Pellman, 2012; Burrell et al., 2013) and less is known if defective mitotic control directly induces chromosomal bridges.

The ultimate purpose of mitosis is to segregate the replicated chromatid pairs and convey the genome to daughter cells completely. The final stage of sister chromatid separation proceeds during the metaphase-to-anaphase transition, which is driven primarily by the activity of the ubiquitin ligase called anaphase-promoting complex, or cyclosome (APC/C). Polyubiquitination and degradation of the substrate proteins, securin and cyclin B1, induces activation of the cohesin-severing protease separase (Reviewed in Kamenz and Hauf, 2017). The spindle assembly checkpoint originating from unattached kinetochores is an inhibitory signal for the APC/C and serves as a surveillance mechanism for sister chromatid separation (Reviewed in Musacchio, 2015). Thus, to eventually induce separase activation, a series of biochemical reactions take place during the metaphase-to-anaphase transition. Vigorous regulation of these processes must underlie mitotic fidelity; however it remains unclear if and how defects in the regulation may contribute to the genomic instability in cancers.

Separase overexpression has been found in different cancer types and shown to have oncogenic activity in mouse models (Zhang et al., 2008; Mukherjee et al., 2014), placing separase as a potential therapeutic target. In this study, by using a probe specific for separase activity, we examined the activation kinetics of separase in live cells and found that cancer cells are widely deficient in its characteristic robust regulation. Our data indicated that prolonged metaphase causally affects activation kinetics of separase, which culminates in chromosome nondisjunction in anaphase.

## Results

To characterize the activation profile of separase in cancer, we performed live cell imaging analysis and measured its activity using the separase probe for various types of cell lines. As previously described, the reporter construct contains a polypeptide including separase-cleavage sites of Scc1, and fused with green (EGFP) and red (mCherry) fluorophores at both ends (Fig. 1A; Shindo et al., 2012). This was located on chromosomes by tethering H2B to the mCherry end, and thus separase-mediated cleavage of Scc1 peptide can be detected by the release of EGFP signal from mCherry on chromosomes (Supplementary Fig. 1). The probe was introduced in principle by plasmid transfection, but for those unsuitable for transfection experiments we adopted lentivirus vector.

**Figure 1.**
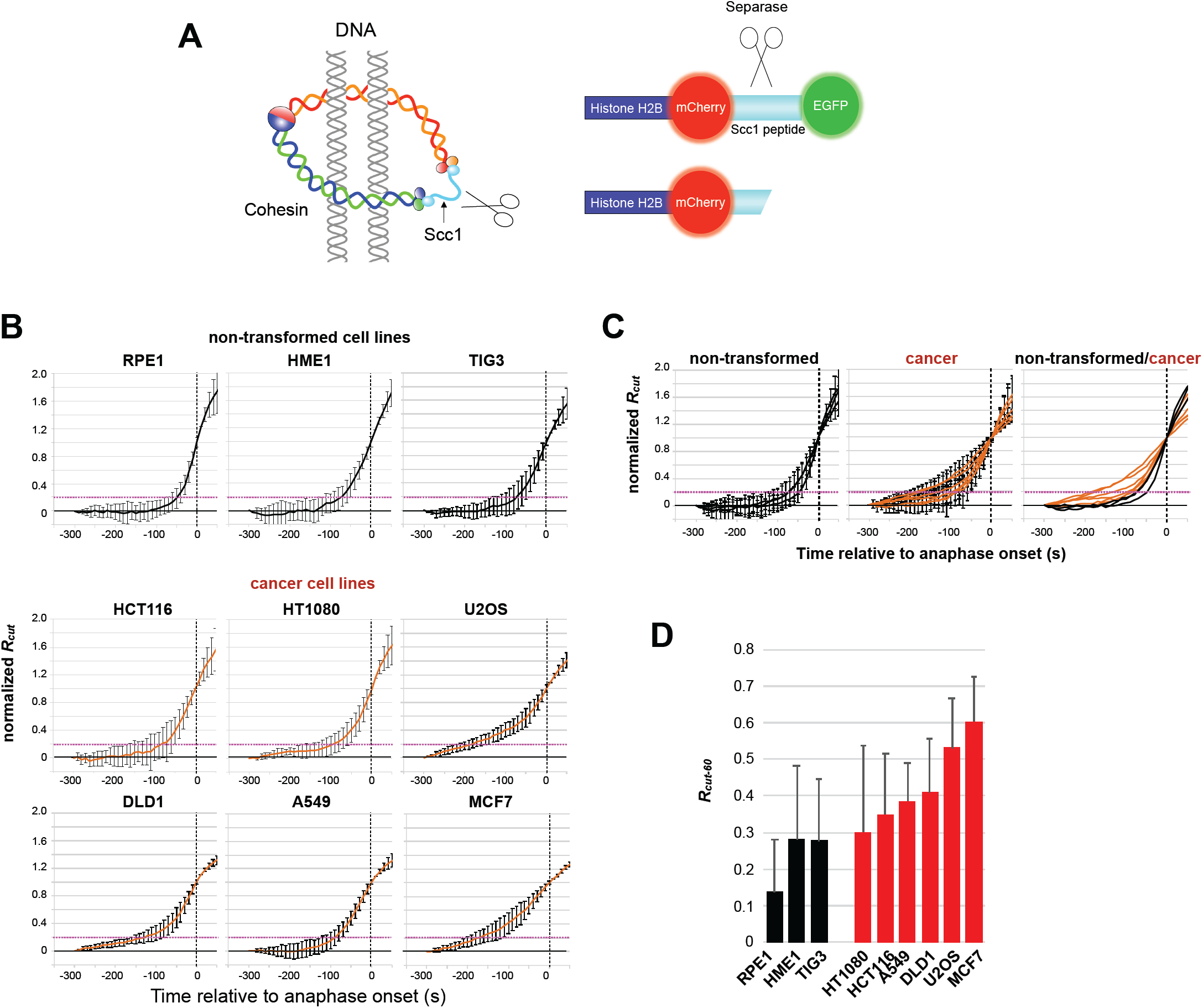
Separase activation profiles in non-transformed diploid and cancer cells. (A) The probe for separase activity. The amount of EGFP fluorophore freed from chromosomes reflects the separase activity. (B) Normalized *R*_*cut*_ values, reflecting the cumulative ratio of cleaved probe, from non-transformed diploid (RPE1, HME1 and TIG3) and cancer (HCT116, HT1080, U2-OS, DLD1, A549 and MCF7) cells were averaged over multiple cells and plotted in black and orange, respectively (n≧10, mean ± SD). Dotted lines in magenta indicate normalized *R*_*cut*_ values at 0.2. (C) Graphs in (B) were combined for non-transformed cell lines and/or cancer cell lines, as indicated. (D) *R*_*cut-60*_, normalized *R*_*cut*_ values at timepoint -60 s, measured in each cell lines were shown in histogram. Error bars are SD.

In non-transformed, chromosomally stable diploid cell lines, such as RPE1, HME1 and TIG3, separase revealed a characteristic robust activation at the metaphase-to-anaphase transition. To quantitatively analyze the levels of fluorescence signals, we calculated a parameter *R*_*cut*_ for each timeframe, reflecting the cumulative ratio of cleaved Scc1 peptide mediated by active separase. The resulting data indicated that the activity of separase is strictly suppressed during much of metaphase until it surges shortly before anaphase onset. By contrast, in cancer cell lines, the activation of separase became detectable already far before the anaphase onset, which was then followed by a gradual increase of its activity until when its main wave of activation eventually induces chromosome separation in anaphase.

Setting a provisional threshold for *R*_*cut*_ at 0.2, which indicates that 20% of the reporter peptide has been cleaved where 100% indicates the ratio reached at the onset of anaphase (Fig. 1B, line in magenta), non-transformed cells surpassed this threshold 60 sec before the anaphase onset (−60 sec). However, in cancer cell lines, typically in DLD1, U2OS and MCF7 cells, this threshold level was exceeded already 120 sec before anaphase onset, reflecting their earlier activation of separase (Fig. 1B, C). Alternatively, when *R*_*cut*_ at timepoint -60 sec, *R*_*cut-60*_, was compared, the proportion of the cleaved peptide ranged between 30 to 60% in cancer cell lines when it was still 12∼28% in non-transformed cell lines (Fig. 1D). These results suggest that cancer cells fail to suppress the activity of separase through metaphase and thus allow its activation at earlier times with regard to anaphase onset.

To ask if this precocious activation is a trait widely shared among cancer cells, instead of cell line-specific phenomena, we examined separase activation profile of RPE1 cells in its transformed line, which was generated by manipulating oncogenes and tumor suppressors (Shirnekhi et al., 2017; Abe et al., 2016). Examination of these otherwise genetically identical two cell lines, we found that the transformed RPE1 cells lacked robustness of separase regulation, as found in various types of cancer cell lines (Fig. 2A, Supplementary Fig. 2). The *R*_*cut-60*_ value resulted in 12% and 35% in parental and transformed cells, respectively. These results indicated that kinetics of separase activation is affected during cellular transformation (Fig. 2B).

**Figure 2.**
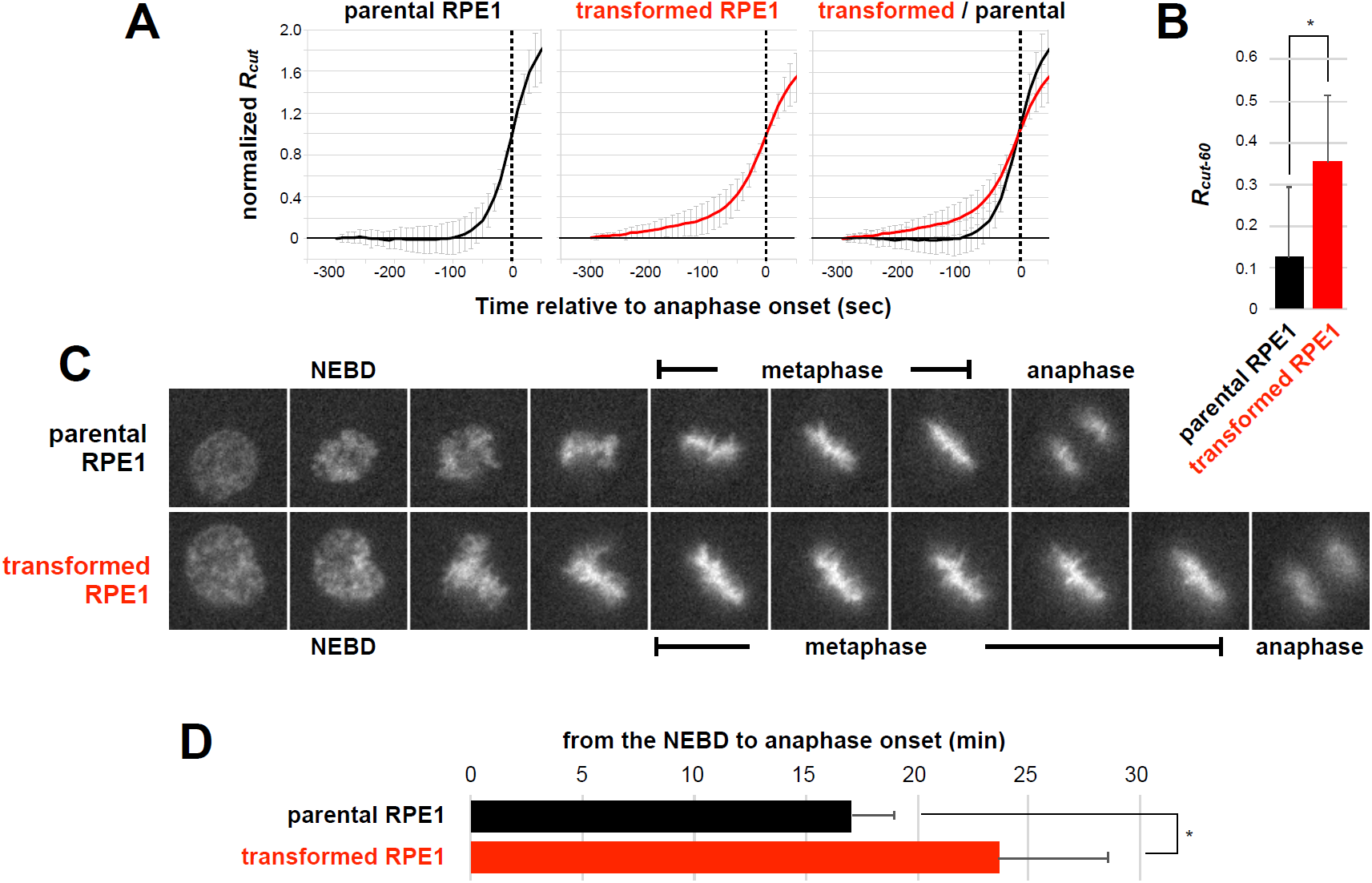
Deregulation of separase in transformed RPE1 cells. (A) Artificially transformed RPE-1 cells (Abe et al., 2016) and its parental RPE-1 cells were analyzed by the separase probe and normalized *R*_*cut*_ was plotted in red and black, respectively (mean ± SD). (B) The *R*_*cut-60*_ values of parental and transformed RPE-1 cell lines were shown in the histogram. Error bars are SD. (C and D) Mitotic length of parental and transformed RPE1 cell lines was analyzed and determined as 17.0 ± 1.9 min and 23.7 ± 4.9 min, respectively (mean ± SD, n=23).*p < 0.0001, two-tailed Student’s t test.

In these live imaging analyses, we noticed that transformed RPE1 cells spent longer times in metaphase (Fig. 2C), which caused prolonged mitosis, from the NEBD to anaphase onset, by approximately 6 min on average than its parental RPE1 cells (Fig. 2D). It has been pointed out that cancer cells are widely associated with prolonged mitosis, with extended metaphase (Therman et al., 1984) and are defective in releasing from mitotic checkpoint arrest (Yang et al., 2008). Consistent with these studies, most cancer cell lines we examined revealed prolonged mitosis (Supplementary Fig. 3).

Intrigued by this common feature of cancer and transformed cell lines, we hypothesized that the tardy progression through metaphase, i.e., delaying anaphase onset, might affect separase regulation. To test this idea, we challenged cells with a low dose of microtubule poison nocodazole, at the concentration that does not perturb overall spindle assembly and allows mitotic progression but causes metaphase retardation. We found 15 ng/ml of nocodazole has such effects to the RPE1 cells; the treatment caused an extension of metaphase by ∼6 min on average, reaching to the range of metaphase length in transformed RPE1 cells (Fig. 3A). Under these conditions, the separase probe indicated that separase undergoes precocious activation following prolonged metaphase, in a similar manner to cancer and transformed cell lines (Fig. 3B). The measurement of their *R*_*cut-60*_ value (∼35%) indicated that the activity of separase failed to be suppressed during prolonged metaphase (Fig. 3C).

**Figure 3.**
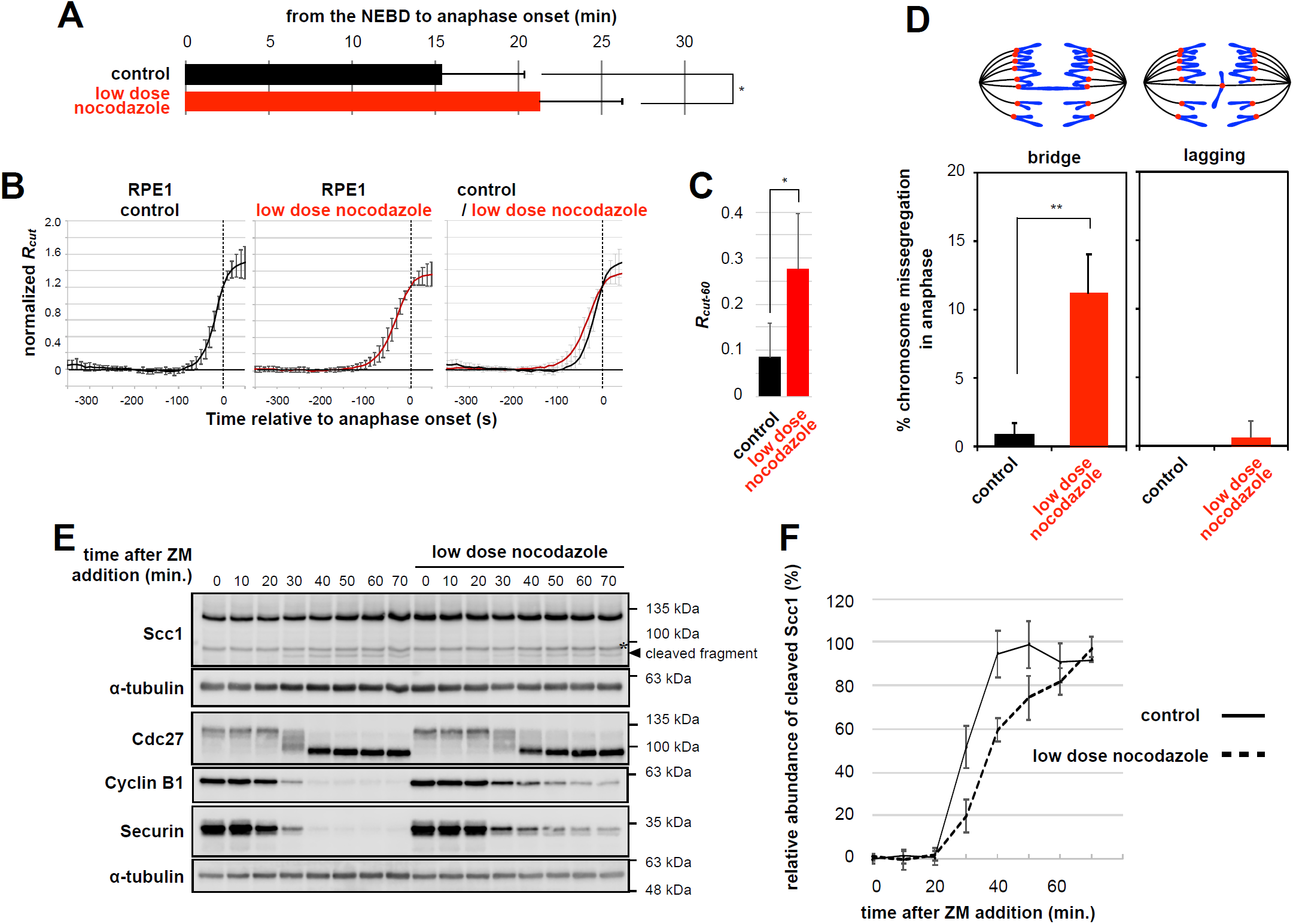
Separase deregulation by delaying anaphase onset. (A) RPE1 cells were treated with 15 ng/mL nocodazole and mitotic length was measured in live cell imaging analysis. Mitotic length of cells in the absence or presence of low dose nocodazole treatment were determined as 15.4 ± 2.4 min and 21.3 ± 4.8 min, respectively (mean ± SD, n>100). (B) Normalized *R*_*cut*_ values from non-treated and 15 ng/mL nocodazole treated RPE-1 cells were plotted in black and red, respectively (mean ± SD). (C) The *R*_*cut-60*_ values were shown in histogram (mean ± SD, n>10).*p <0.0001, two-tailed Student’s t test. (D) Chromosome missegregation caused by low dose nocodazole treatment is mostly chromosomal bridges, instead of lagging chromsomes. RPE1 cells were fixed and stained with DAPI to visualize chromosomes after treating with or without 15 ng/ml nocodazole. Bar graph indicates mean ± SD from three independent experiments. ** p < 0.02, two-tailed Student’s t test.(E) HeLa (Kyoto) cells were treated with 7.5 μM *S*-Trityl-L-cysteine for 18 h and mitotic cells were collected and treated with 5 μM ZM447439 either with or without 10 ng/mL nocodazole for the time indicated. Cells were harvested and subjected to immunoblot analysis. (F) Kinetics of cohesin cleavage. Intensities of Scc1 cleaved fragment (arrowhead in E) were measured and plotted over time after subtracting the minimum value from all data points and divided by the maximum value to set the point for 100 % cleavage (mean ± SD).

Importantly, in the following anaphase, the nocodazole-treated RPE1 cells revealed an increasing rate of chromosome missegregation (Fig. 3D). To our surprise, the majority of these were judged as chromosomal bridges (11.2±2.9%), instead of lagging chromosomes (0.6±1.1%). Chromosomal bridges are thought to stem from sister chromatid dissociation defects, the origins of which may include defective chromatid assembly, incomplete replication, and dicentric chromosomes being pulled apart (Ganem and Pellman, 2012). Provided that deregulation of separase causally relates to chromosomal bridges, a plausible possibility was that the bridges observed in our experiment is due to the insufficient removal of cohesin at anaphase onset.

To address this possibility, we asked if separase-mediated cohesin cleavage is affected by perturbing the robust activation of separase with nocodazole. To do this, we prepared mitotic cell population and released from the arrest to let cells traverse from metaphase to anaphase in a highly synchronous manner (Shindo et al., 2012). When the cells were released in the absence of nocodazole, the cleaved fragment of cohesin subunit Scc1 appeared at the expected molecular weight, about 90 kDa, at 30 min after the release and topped off in next 10 min (Fig. 3E and F). When the cells were released in the presence of low dose nocodazole, degradation of securin and cyclin B1 was delayed reflecting the retarded metaphase progression. The fragment can be hardly detected at 30 min and gradually increased its intensity and apparently lower through the reasonable period of experiment (Fig. 3E and F), indicating that total amount of cleaved Scc1 at the onset of anaphase would be much lower in prolonged mitosis. These results support the interpretation that anaphase bridges are caused by incomplete removal of cohesin from chromosomes, and that the robust activation kinetics of separase is required to attain sufficient level of its activity.

These results are consistent with the notion that a delay in metaphase progression perturbs the robust activation of separase, which seemed essential to gain sufficient levels of its output to sever sister chromatids. Finally, we reasoned that the profile of separase activity and the intact chromosome segregation might restore, if slow metaphase progression can regain its kinetics. To test this possibility, we challenged cells with an Mps1 inhibitor reversine to facilitate inactivation of the mitotic checkpoint, as metaphase is driven by checkpoint inactivation (Santaguida et al., 2010). Treating transformed RPE1 cells with 125 nM of reversine shortened their prolonged metaphase up to the length normally seen in parental RPE1 cells (Figure 4A). Under these conditions, we found that cells restored the robust kinetics of separase activation (Figure 4B and C), and concomitantly, the rate of chromosomal bridges significantly declined from 15.6% to 6.7% (Figure 4D). The rate of lagging chromosomes increased from 3.6% to 8.7%, probably due to remaining erroneous kinetochore-microtubule attachments that would have been corrected during prolonged metaphase in these transformed cells. Based on these results we concluded that the progression of metaphase directly affects both activation and functional profile of separase.

**Figure 4.**
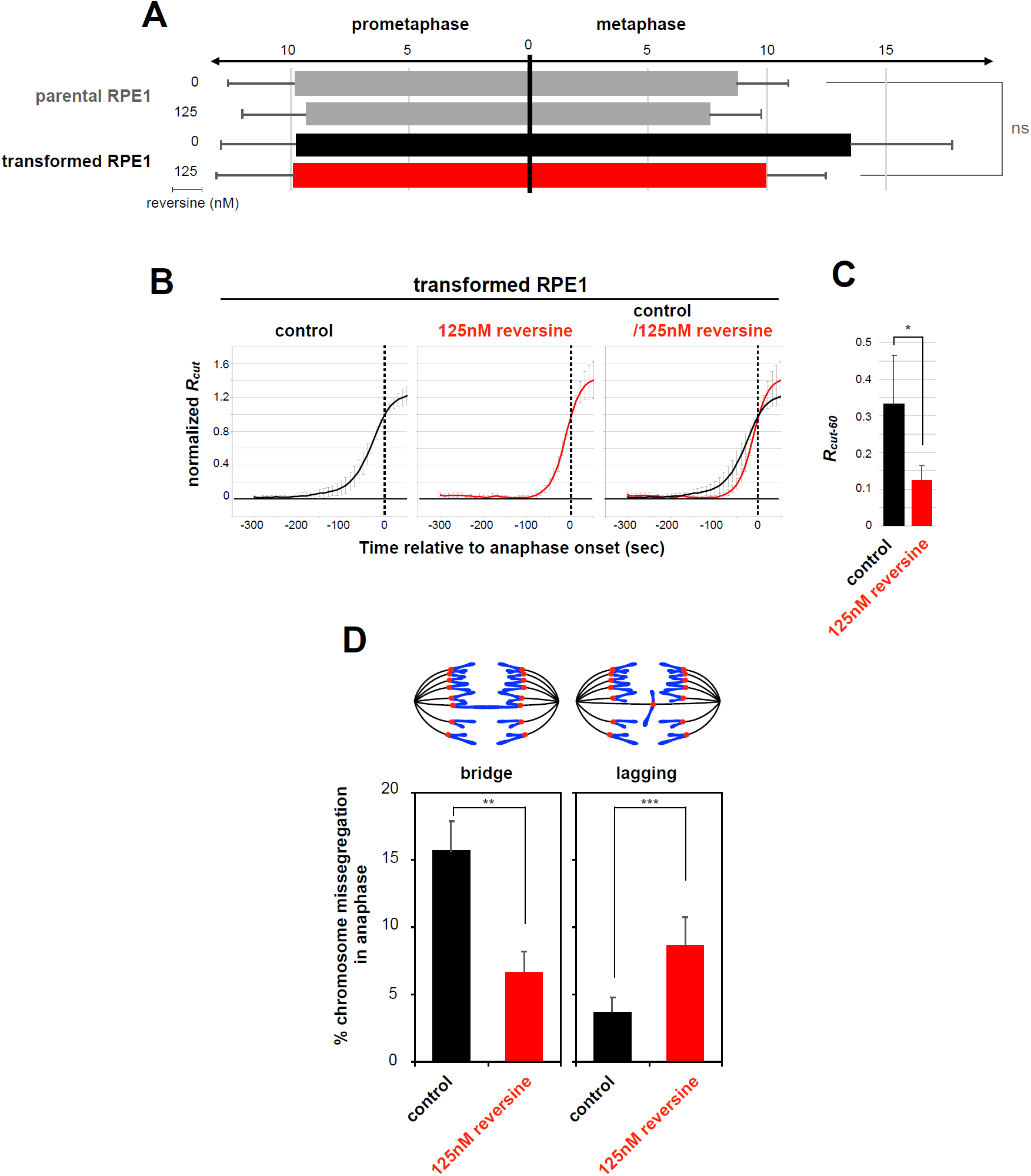
Restoration of the robust separase regulation and function by shortening prolonged metaphase. (A) Parental and transformed RPE1 cells were treated with or without 125 nM reversine (Santa cruz) and mitotic lengths were measured in live cell imaging analysis. Reversine treatment shortened mitotic lengths from 18.7 ± 3.4 min to 17.0 ± 2.8 min in parental cells, and from 23.2 ± 5.4 min to 19.9 ± 3.6 min in transformed cells, respectively. The onset of metaphase was determined when all chromosomes aligned to the metaphase plate. Reversine treatment shortened metaphase length from 8.7± 2.1 min to 7.6 ± 2.1 min in parental cells, and from 13.5± 4.3 min to 10.0 ± 2.5 min in transformed cells, respectively (mean ± SD, n>70). Note that lengths of metaphase largely account for shortening total mitotic lengths in this experiment. (B) Normalized *R*_*cut*_ values from untreated and 125 nM reversine treated transformed RPE-1 cells were plotted in black and orange, respectively (mean ± SD, n>10). (C) The *R*_*cut-60*_ values were shown in histogram (mean ± SD, n>10). (D) Transformed RPE1 cells were fixed and stained with DAPI to visualize chromosomes, after treating with or without 125 nM reversine. Bar graph indicates mean ± SD from three independent experiments. \**p* < 0.0001, ** p < 0.01, ***p < 0.05, two-tailed Student’s t test.

## Discussion

The probe for separase indicates its spatiotemporal regulation on a single cell basis, which allowed us to find that cancer cells have widely altered the activation kinetics: separase activates precociously during prolonged metaphase. Remarkably, this loss of robust separase activation culminated in an insufficient removal of cohesin by anaphase onset, manifested by the emergence of chromosomal bridges. These observations let us to propose a previously unanticipated etiology for chromosome instability. The observations in cancers in return indicate that a swift and decisive transition from metaphase-to-anaphase is intrinsic to the mitotic control, and importantly this comprises an essential cellular function to prevent pathological segregation errors.

Anaphase is thought to be the most critical and vulnerable period in the cell cycle in propagating of the genome (e.g., Bizard and Hickson, 2018). To which extent might this continued chromosome nondisjunction events following defective separase regulation contribute to the genomic instability observed in human tumors is a following important question. Identification of the breakage sites or genomic loci associated with separase deregulation should be informative. Arresting cells for hours in mitosis, i.e., mitotic arrest, is also known to induce various types of DNA damage (Reviewed in Ganem and Pellman, 2012). It should be noted however that prolonged metaphase observed in cancer cells is only at most several tens of minutes, and thus mitotic arrest must evoke different problems from what we pointed out in this study.

What accounts for prolonged metaphase in cancers? Given that strength of the spindle-assembly checkpoint signaling correlates with the number of unattached kinetochores (Dick and Gerlich, 2013; Collin et al., 2013), cancer cells with increased number of chromosomes (hyperploid) will require longer time for the attachment and thereby delays the checkpoint inactivation process. It is also known that spindle microtubules are less dynamic (more stable) in cancer cells (Bakhoum et al., 2009a) and this property must affect the efficiency of chromosomes to be attached to spindle microtubules. In addition, they often accompany supernumerary centrosomes and develop multipolar spindle in mitosis, which would also require additional times to be corrected (Silkworth et al., 2009; Ganem et al., 2009). These circumstances will affect the checkpoint inactivation processes and delay metaphase progression.

Because it is during prolonged metaphase when separase begin to activate precociously, there seems at least two conditions to attain the robust activation kinetics. One is that the spindle-assembly checkpoint needs to be switched off promptly as soon as it is satisfied, so that cells do not spend longer times in metaphase. The other is that separase molecules must restrain sporadic sparks until they become active more or less simultaneously, in order to yield the primary activation wave. Is there a causative link between these two sequential events? As the retarded checkpoint inactivation inversely decelerates the APC/C activation kinetics and degradation of separase inhibitors, securin and cyclin B1 (Uchida et al., 2009), one prediction is that a slower decline of separase inhibitors leads to the state allowing separase activation before the right time.

Furthermore, the precocious activation is associated with lower output of separase activity and failed to clear all the cohesive cohesin between sister chromatids (Figure 3), underscoring the relevance of the synchronicity of separase activation. We can speculate two scenarios explaining why precocious activation attenuates the net separase activity at the onset of anaphase. First, the activation process is irreversible and allows separase to spark only once in a short period of time. The precociously activated separase will not contribute to the primary activation wave preceding anaphase onset and the total amount of active separase becomes smaller. Supporting this idea, the activated form of separase is found to be short lived (Hellmuth et al., 2015). A second, more challenging idea is that, separase activity is propagated through an *in trans* activation, as in the case for apoptosis inducing protease caspase, and thus asynchronous activation is disadvantageous in conveying the activity. Presence of already activated protease would act as a decoy in the *in trans* activation mechanism. Testing these hypotheses in future works will provide a clue to understanding mechanistically how separase achieves the robust activation.

In conclusion, our results show that a wide range of cancer cells consistently have an impaired regulation in releasing the spindle-assembly checkpoint ‘brake’ and subsequent activation of separase ‘accelerator’, which results in chromosomal bridges in anaphase. These findings significantly imply that a swift release of the brake and a rapid ignition of the engine are essential cellular function to drive through the metaphase-to-anaphase safely.

## Acknowledgments

We are grateful to Jennifer DeLuca and Jacob Herman for sharing the transformed RPE1 line; Utako Kato and Youko Hirayama for technical assistance; and all members of the T.H. laboratory for discussions. Research in the T.H. lab is supported by the Japan Society for the Promotion of Science (JSPS) Grant-in-Aid for Scientific Research (15H02365, 18H04034, 15H05977 [to T.H.] and 25711003, 19K07677 [to N.S.]).

## Materials and methods

### Antibodies

Rabbit polyclonal antibody to Scc1(#8277) was raised against two synthetic peptides DEPIIEEPSR (corresponding to amino acid sequence number 441-450) and ATPGPRFHII (622-631). The following mouse monoclonal antibodies were used: Separase (XJ11-1B12, K0201-3; MBL, Nagoya, Japan), Cdc27 (35/CDC27, 610455, BD Biosciences), Cyclin B1 (GNS1, Santa Cruz), Securin (EPR3240, ab79546, Abcam), alpha-tubulin (B512, T6074, Sigma).

### Cell culture

HeLa Kyoto (Gift from S. Narumiya, Kyoto University), hTERT-RPE-1 (RPE-1, purchased from ATCC), TIG3, HCT116, HT1080, U2OS, A549 and MCF7 were maintained in Dulbecco’s modified Eagle’s medium (DMEM). HME1 was maintained in HMEGM (LONZA) without GA-1000 and DLD1 was maintained in RPMI1640 medium. In all cases, medium was supplemented with 10% fetal bovine serum (FBS) and cells were maintained at 37 °C in an atmosphere of 5% CO_2_.

### Quantitative analysis of separase activity

H2B-mCherry-Scc1(142-175aa)-EGFP from original separase sensor construct (Shindo et al., 2012) was subcloned into the Kpn1 and NotI sites of lentiviral vector pLenti-Easy-HA. Recombinant lentivirus was produced by transient transfection of 293T cells using Virapower packaging mix (Invitrogen). Cells were plated into 2 well LabTek chambered coverglass (Nunc) and transduced with the lentivirus carrying separase sensor. After 24 h, cells were washed twice with PBS and medium was changed to Leibovitz’s L-15 medium (GIBCO) supplemented with 20% fetal bovine serum, 20 mM HEPES (pH7.6) for live cell imaging. Sensor expressing metaphase cells were chosen and images were captured every 10 s for up to 1 h with a 100x objective. Obtained images were analyzed using ImageJ software. Regions of chromosomes were determined using mCherry signals by thresholding and the mean value of EGFP and mCherry signals of these defined region was calculated using Image Calculator plugin by multiplying each image by the thresholded mCherry image. Resulting values were normalized to the value at time point -300 s before anaphase onset (I^EGFP^ and I^mCherry^, respectively) and R_cut_ = 1- (I^EGFP^ / ImCherry) value from each time point was calculated for each cell analyzed. Because expression levels of separase sensor vary between the cells, R_cut_ values were normalized to the R_cut_ values at time point 0 s (anaphase onset) and average of that values were plotted. The normalized R_cut_ value stands for the relative ratio of cleaved sensor as compared to the anaphase onset and R_cut-60_ (R_cut_ at -60 s, see text) was used to assess precocious activation of separase. For live imaging of RPE-1 cells, the medium was supplemented with 10 ng/mL Hoechst 33342 to visualize DNA as lentivirus transduced RPE-1 cells had unknown mCherry signals apart from chromosomes which hindered the following ImageJ analyses.

### Live cell imaging analysis

Cells were plated into Leibovitz’s L-15 medium (GIBCO) supplemented with 20% fetal bovine serum, 20 mM HEPES (pH7.6) and 10 ng/mL Hoechst 33342 on LabTek chambered coverglass (Nunc), and the chamber lids were sealed with silicone grease. Images were captured every 3 min for up to 12 h with a 20x objective.

### Cell synchronization

Synchronous cell population traversing through the metaphase to anaphase was obtained based on the modified procedure described in Shindo et al., (Shindo et al., 2012). Logarithmically proliferating HeLa Kyoto cells were treated with Eg5 inhibitor 7.5 μM *S*-Trityl-L-cysteine (Tocris Biosciences) for 18 h. Mitotic cells were then collected and treated with 5 μM ZM447439 (Tocris Biosciences) either with or without 10 ng/mL nocodazole for the time stated before being harvested for immunoblot analyses.

---

**Supplementary Figure 1.**
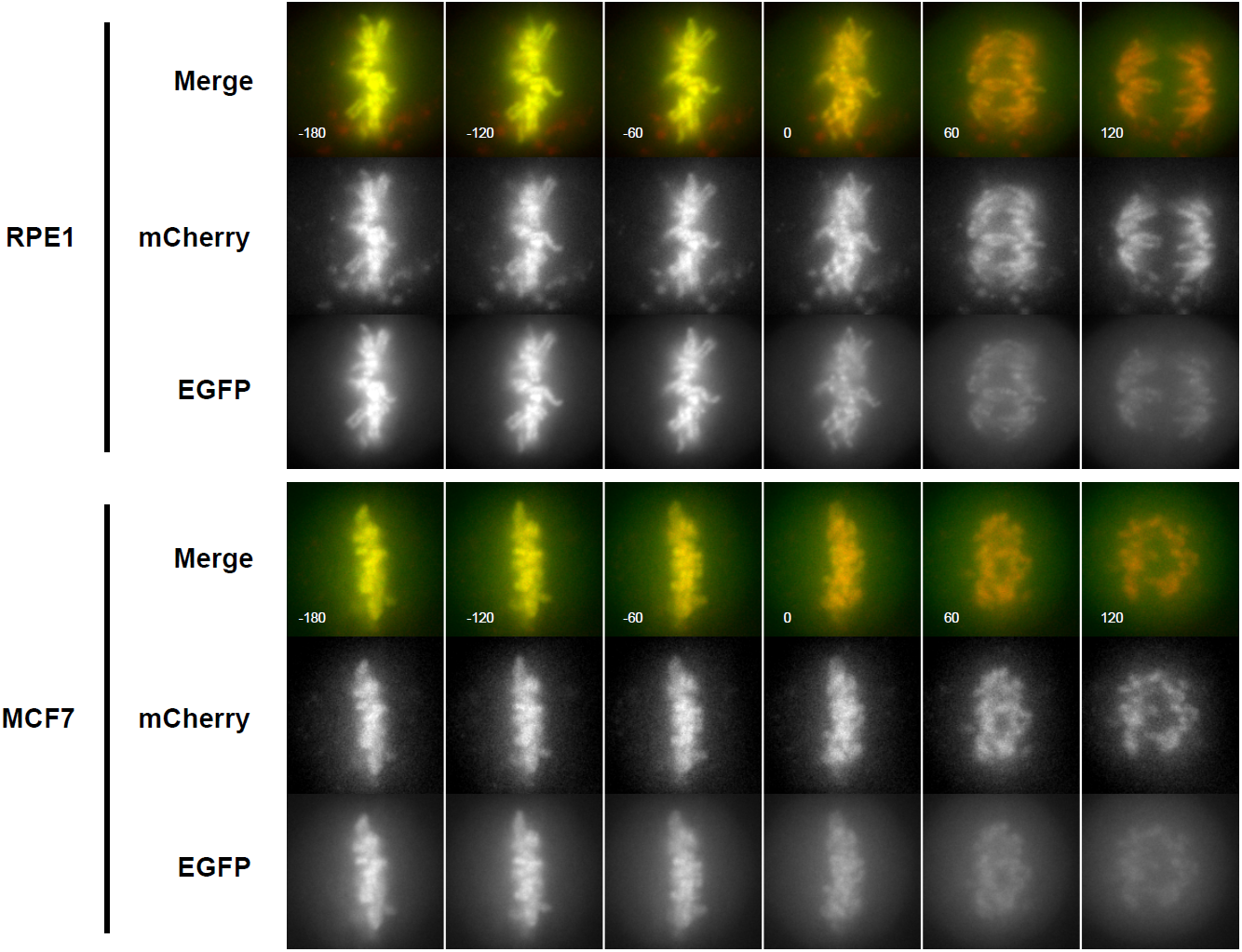
Quantification of separase activity in non-transformed and cancer cells. Still images extracted from live-cell imaging experiments were aligned on the time axis (every 60 s) according to anaphase onset as determined by sister chromatid separation. Representative images from RPE1 and MCF7 cells were shown. Note that cytoplasmic EGFP signals can be detected from earlier timepoints in MCF7 than in RPE1 cells.

**Supplementary Figure 2.**
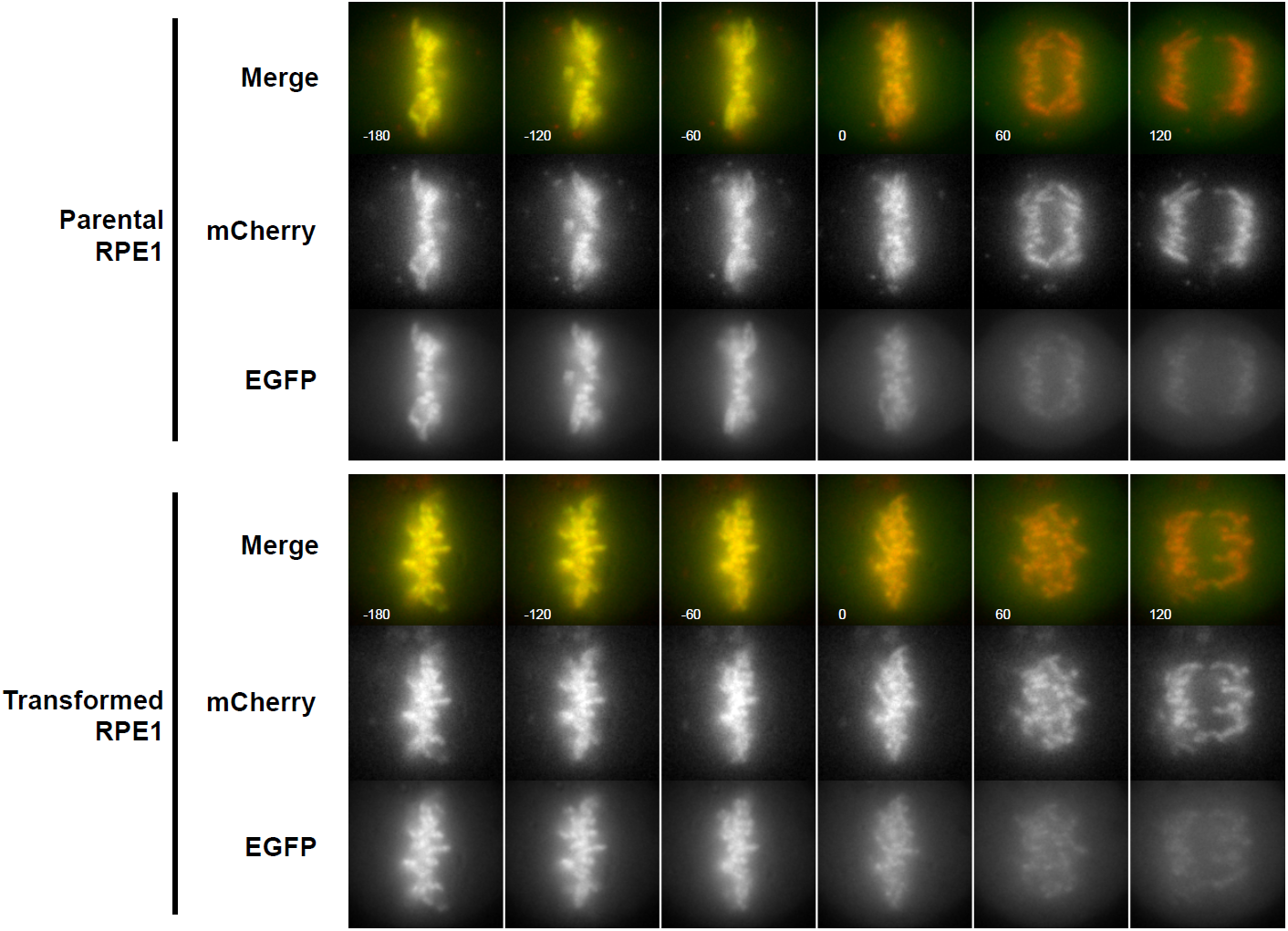
Quantification of separase activity in transformed RPE1 and its parental non-transformed RPE1cells. Still images extracted from live-cell imaging experiments were aligned on the time axis (every 60 s) according to anaphase onset as determined by sister chromatid separation. Representative images from transformed and untransformed RPE1 cells were shown.

**Supplementary Figure 3.**
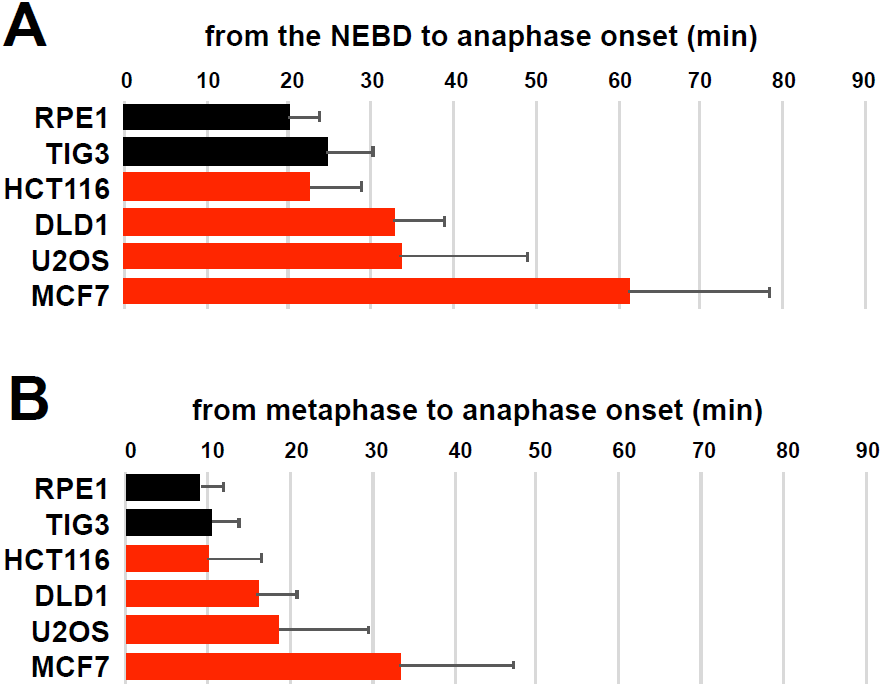
Prolonged mitosis in cancer cell lines. Mitotic cells were extracted from a long term live cell imaging analysis of asynchronous culture, and the time from the NEBD to anaphase onset (A) and the time from metaphase alignment to anaphase onset, i.e., length of metaphase (B) were summarized in the histogram for indicated cell lines (mean ± SD, n>55).

## Notes

### Competing Interest Statement

The authors have declared no competing interest.

## References

Abe, Y., Sako, K., Takagaki, K., Hirayama, Y., Uchida, K.S.K., Herman, J.A., Deluca, J.G., Hirota, T., 2016. HP1-Assisted Aurora B Kinase Activity Prevents Chromosome Segregation Errors. Dev Cell 36, 487–497. doi:10.1016/j.devcel.2016.02.008

Bakhoum, S.F., Genovese, G., Compton, D.A., 2009a. Deviant kinetochore microtubule dynamics underlie chromosomal instability. Curr Biol 19, 1937–1942. doi:10.1016/j.cub.2009.09.055

Bakhoum, S.F., Thompson, S.L., Manning, A.L., Compton, D.A., 2009b. Genome stability is ensured by temporal control of kinetochore-microtubule dynamics. Nat Cell Biol 11, 27–35. doi:10.1038/ncb1809

Bizard, A.H., Hickson, I.D., 2018. Anaphase: a fortune-teller of genomic instability. Curr Opin Cell Biol 52, 112–119. doi:10.1016/j.ceb.2018.02.012

Burrell, R.A., McClelland, S.E., Endesfelder, D., Groth, P., Weller, M.-C., Shaikh, N., Domingo, E., Kanu, N., Dewhurst, S.M., Gronroos, E., Chew, S.K., Rowan, A.J., Schenk, A., Sheffer, M., Howell, M., Kschischo, M., Behrens, A., Helleday, T., Bartek, J., Tomlinson, I.P., Swanton, C., 2013. Replication stress links structural and numerical cancer chromosomal instability. Nature 494, 492–496. doi:10.1038/nature11935

Cimini, D., Wan, X., Hirel, C.B., Salmon, E.D., 2006. Aurora kinase promotes turnover of kinetochore microtubules to reduce chromosome segregation errors. Curr Biol 16, 1711–1718. doi:10.1016/j.cub.2006.07.022

Collin, P., Nashchekina, O., Walker, R., Pines, J., 2013. The spindle assembly checkpoint works like a rheostat rather than a toggle switch. Nat Cell Biol 15, 1378–1385. doi:10.1038/ncb2855

Dick, A.E., Gerlich, D.W., 2013. Kinetic framework of spindle assembly checkpoint signalling. Nat Cell Biol 15, 1370–1377. doi:10.1038/ncb2842

Ganem, N.J., Godinho, S.A., Pellman, D., 2009. A mechanism linking extra centrosomes to chromosomal instability. Nature 460, 278–282. doi:10.1038/nature08136

Ganem, N.J., Pellman, D., 2012. Linking abnormal mitosis to the acquisition of DNA damage. 199, 871–881. doi:10.1083/jcb.201210040

Hellmuth, S., Rata, S., Brown, A., Heidmann, S., Novak, B., Stemmann, O., 2015. Human Chromosome Segregation Involves Multi-Layered Regulation of Separase by the Peptidyl-Prolyl-Isomerase Pin1. Mol Cell 58, 495–506. doi:10.1016/j.molcel.2015.03.025

Kamenz, J., Hauf, S., 2017. Time To Split Up: Dynamics of Chromosome Separation. Trends Cell Biol 27, 42–54. doi:10.1016/j.tcb.2016.07.008

Mankouri, H.W., Huttner, D., Hickson, I.D., 2013. How unfinished business from S-phase affects mitosis and beyond. EMBO J 32, 2661–2671. doi:10.1038/emboj.2013.211

Mukherjee, M., Ge, G., Zhang, N., Edwards, D.G., Sumazin, P., Sharan, S.K., Rao, P.H., Medina, D., Pati, D., 2014. MMTV-Espl1 transgenic mice develop aneuploid, estrogen receptor alpha (ERα)-positive mammary adenocarcinomas. Oncogene 33, 5511–5522. doi:10.1038/onc.2013.493

Musacchio, A., 2015. The Molecular Biology of Spindle Assembly Checkpoint Signaling Dynamics. Curr Biol 25, R1002–18. doi:10.1016/j.cub.2015.08.051

Santaguida, S., Tighe, A., D’alise, A.M., Taylor, S.S., Musacchio, A., 2010. Dissecting the role of MPS1 in chromosome biorientation and the spindle checkpoint through the small molecule inhibitor reversine 190, 73–87. doi:10.1083/jcb.201001036

Shindo, N., Kumada, K., Hirota, T., 2012. Separase sensor reveals dual roles for separase coordinating cohesin cleavage and cdk1 inhibition. Dev Cell 23, 112–123. doi:10.1016/j.devcel.2012.06.015

Shirnekhi, H.K., Kelley, E.P., Deluca, J.G., Herman, J.A., 2017. Spindle assembly checkpoint signaling and sister chromatid cohesion are disrupted by HPV E6-mediated transformation. Mol Biol Cell 28, 2035–2041. doi:10.1091/mbc.E16-12-0853

Silkworth, W.T., Nardi, I.K., Scholl, L.M., Cimini, D., 2009. Multipolar spindle pole coalescence is a major source of kinetochore mis-attachment and chromosome mis-segregation in cancer cells. PLoS ONE 4, e6564. doi:10.1371/journal.pone.0006564

Therman, E., Buchler, D.A., Nieminen, U., Timonen, S., 1984. Mitotic modifications and aberrations in human cervical cancer. Cancer Genet. Cytogenet. 11, 185–197. doi:10.1016/0165-4608(84)90113-4Get

Thompson, S.L., Bakhoum, S.F., Compton, D.A., 2010. Mechanisms of chromosomal instability. Curr Biol 20, R285–95. doi:10.1016/j.cub.2010.01.034

Uchida, K.S.K., Takagaki, K., Kumada, K., Hirayama, Y., Noda, T., Hirota, T., 2009. Kinetochore stretching inactivates the spindle assembly checkpoint 184, 383–390. doi:10.1083/jcb.200811028

Yang, Z., Loncarek, J., Khodjakov, A., Rieder, C.L., 2008. Extra centrosomes and/or chromosomes prolong mitosis in human cells. Nat Cell Biol 10, 748–751. doi:10.1038/ncb1738

Zhang, N., Ge, G., Meyer, R., Sethi, S., Basu, D., Pradhan, S., Zhao, Y.-J., Li, X.-N., Cai, W.-W., El-Naggar, A.K., Baladandayuthapani, V., Kittrell, F.S., Rao, P.H., Medina, D., Pati, D., 2008. Overexpression of Separase induces aneuploidy and mammary tumorigenesis. Proceedings of the National Academy of Sciences 105, 13033–13038. doi:10.1073/pnas.0801610105

